# Rapamycin as a potent and selective inhibitor of vascular endothelial growth factor receptor in breast carcinoma

**DOI:** 10.1101/2020.08.27.269688

**Authors:** Muhammad Shahidan Muhammad Sakri, Wan Faiziah Wan Abdul Rahman, Tengku Ahmad Damitri Al-Astani Tengku Din, Hasnan Jaafar, Vinod Gopalan

## Abstract

Angiogenesis is the process of new vascular formation, which is derived from various factors. For suppressing cancer cell growth, targeting angiogenesis is one of the therapeutic approaches. Vascular endothelial growth factor family receptors, including Flt-1, Flk-1, and Flt-4, have been found to play an essential role in regulating angiogenesis. In the present study, we evaluated the effects of rapamycin and platelet factor-4 toward breast carcinoma at the proteomic and genomic levels. A total of 60 N-Methyl-N-Nitrosourea-induced rat breast carcinomas were treated with rapamycin, platelet factor-4, and rapamycin+platelet factor-4. The tumors were subsequently subjected to immunohistological protein analysis and polymerase chain reaction gene analysis. Protein analysis was performed using a semi-quantitative scoring method, while the mRNA expression levels were analyzed based on the relative expression ratio. There was a significant difference in the protein and mRNA expression levels for the selected markers. In the rapamycin+platelet factor-4 treated group, the Flt-4 marker was downregulated, whereas there were no differences in the expression levels of other markers, such as Flt-1 and Flk-1. On the other hand, platelet factor-4 did not exhibit a superior angiogenic inhibiting ability in this study. Rapamycin is a potent anti-angiogenic drug; however, platelet factor-4 proved to be a less effective drug of anti-angiogenesis on rat breast carcinoma model.

## Introduction

Angiogenesis, a process by which new vasculature forms, occurs as a result of a variety of factors. Neo-angiogenesis is one of the main processes that allow tumors to grow larger than 2 mm in diameter [1]. Since the discovery of neo-angiogenesis, numerous intensive research studies have been conducted in order to determine the root factors that influence and enhance this process. Targeting angiogenic factors, such as vascular endothelial growth factor (VEGF), platelet-derived growth factor (PDGF), insulin-like growth factor (IGF), fibroblast growth factor (FGF), and their receptors, have potential impact on the development of new strategies to suppress tumor growth, especially at later stages [2].

Fms-like tyrosine 1 (Flt-1), Fms-like kinase 1 (Flk-1), and Fms-like tyrosine 4 (Flt-4) are members of the VEGF receptor family and are known as prognostic markers [3]. In addition, each of these VEGF receptors has been found to play a role in the regulation of neo-vascularization [3]. Moreover, these receptors co-regulate each other’s’ expression to trigger angiogenesis. Among these, Flk-1, known as a kinase-derived receptor (KDR), plays a significant role in the regulation of angiogenesis, whereas other molecules in the same family support this process [4]. Flt-1 has also been reported to play a significant role in the branching of new arteries, whereas Flk-1 influences vein formation, and Flt-4 plays a vital role in the regulation of lymphatic vessel formation.

Flt-1 is known to be the cognate receptor for VEGF-A. This receptor plays a crucial role in the regulation of neo-angiogenesis alongside Flk-1. Recent studies have revealed that Flt-1 induces angiogenesis along with Flk-1 through several intrinsic pathways, including the PLC-y, Grb2, and PI3K/Akt, which in turn activate downstream pathways, such as MAPK for proliferation, eNOS for cell permeability, and Caspase 9 for survival [5].

Contrarily, Flt-4 plays a significant role in the regulation of lymphangiogenesis, which is a common route of cancer metastases [6]. Most cases of aggressive carcinoma exhibit significant Flt-4 expression, which reflects the degree to which several factors, especially Flt-4, play a role in the regulation of neo-lymphangiogenesis [7]. Based on our previous findings in rats with breast carcinoma, the treated groups exhibited a significant downregulation of the Flt-4 receptor; moreover, the cancer cells were significantly less aggressive, and most of the tumor cells appeared localized. In the present study, we examined and analyzed the expression of VEGFRs in rat breast carcinoma following treatment with rapamycin and PF-4.

## Materials and methods

### Animal procedures

Sixty female Sprague Dawley (SD) rats were obtained from the Animal Research and Services Centre, and ethical clearance was obtained from the Animal Ethics Committee [PPSG/07(A)/044/ (2010) (56)]. The rats were housed at the animal house unit. The caging and handling of the rats conformed to good laboratory practices, as outlined by the Animal Research and Services Centre (ARASC). To avoid discomfort, pain, and stress, the rats were given an analgesic drug (Ketamine) at a dose of 80 mg/kg intraperitoneally during the measurement of the tumor. The rats were maintained in groups of six and fed a standard laboratory diet [8]. They were divided into four groups, and each group was given different interventions/drug(s), as follows: the control (untreated) group 1 [n = 15], rapamycin-treated group 2 [n = 15], platelet factor-4-treated group 3 (PF-4) [n = 15], and rapamycin and PF-4-treated group 4 (rapamycin+PF-4) [n = 15]. The rats were sacrificed 5 days after the drug intervention process.

### NMU, rapamycin, and PF-4 preparation and tumor induction

NMU (N-nitroso-N-methylurea) was dissolved in freshly prepared 0.9% normal saline prior to the induction process. The NMU solution was injected intraperitoneally into 21-day-old rats at a dose of 70 mg/kg body weight [9], after which each animal was monitored and palpated weekly for mammary tumor lesions. Daily inspection and palpation were also performed to monitor the onset of tumors in the mammary region, and the weight of the rats were measured daily. Rapamycin was mixed with absolute ethanol and diluted in mixtures of 10% polyethylene glycol (PEG)-400, 8% ethanol, and 10% Tween-80 to a final concentration of 20 µg/0.2 mL, whereas PF-4 was dissolved in normal saline with a final concentration of 20 µg/0.2 ml

### Study design

This study was conducted *in vivo* and involved an intraperitoneal injection of the NMU carcinogen into 60 female Sprague Dawley rats using a 21G needle to induce breast carcinoma. After the development of breast tumors within 40 days of NMU carcinogen injection, the tumors were then suppressed using an angiogenic inhibitor as either a single treatment (PF-4 or rapamycin) or a combination of these two compounds; for combination treatment, both PF-4 and rapamycin were injected into the tumors. After 5 days of intervention, the rats were terminated, whereas rats in the control group were terminated 45 days after tumor induction. Hematoxylin and eosin staining together with immunohistochemical stains were used to analyze the histology and expression of angiogenic protein markers. All results were then statistically analyzed using the SPSS software version 21.

### Immunohistochemistry

Tissue sections on slides were deparaffinized in xylene and then rehydrated in graded alcohol solutions. Antigen retrieval was conducted in a pressure cooker, and the selection of an appropriate buffer depended on the detected marker. Nonspecific background staining was blocked by a 3% hydrogen peroxide solution. Sections were incubated with the primary antibody for 1 h at room temperature, which was followed with several washes and incubation with the secondary antibody using an UltraVision ONE Large Volume Detection System HRP Polymer Kit. The protein reactivity was visualized using a 3,3′-diaminobenzidine (DAB) Plus substrate system (Cat. No. TL-125-PHJ, LabVision, USA). The tissue was counterstained with hematoxylin, which stained the nuclei.

### Immunohistochemistry scoring system

Each section was scored in a blinded manner by two observers based on a previously reported semiquantitative scoring method [9]. Each sample was scored based on the percentage of tumor cells stained (0-absence of staining; 1-<30% of tumor cells stained; 2-30–60% of tumor cells stained; and 3->60% of tumor cells stained).

### RNA expression analysis

The tissue sample extraction was performed using TRIZOL (Invitrogen, USA) to obtain total RNA. All RNA preparation and handling steps were carried out under RNAse-free conditions and were performed in a laminar flow hood. Total RNA from each fraction was dissolved in 20 μL of RNA storage buffer (Ambion, USA) and stored at −80°C until use. RNA concentration was determined by 1% gel electrophoresis and absorbance readings at 260 nm using the Nanodrop-1000 spectrophotometer (NanoDrop, Technologies, USA). The reading of RNA samples was within the range of 1.8–2.2, indicating the purity of the sample. This ratio was obtained from the 260/280 absorbance ratio. mRNA was isolated from total RNA using the Dynabeads mRNA Purification kit (Invitrogen, USA) according to the manufacturer’s instructions. cDNA synthesis was performed using the SuperScript™ First-Strand Synthesis System (Invitrogen, USA) in a total volume of 20 μL according to the manufacturer’s instructions.

### Quantitative RT-PCR analysis

The expression of 11 genes of interest and three selected reference genes was examined by real-time TaqMan® PCR assay. Expression levels were determined using the exon spanning hydrolysis probes (FAM or MGB dye-labelled), which are commercially available as “Assay on Demand” (Applied Biosystems, Foster City, CA, USA), with optimized primer and probe concentrations as stated in Table 1.

**Table 1.** List of gene markers by TaqMan® PCR assay used for angiogenesis analysis.

| Gene symbol | Assay ID | Gene name |
| --- | --- | --- |
| *Flt-4* | Rn00570815_m1 | fms-related tyrosine kinase 1 |
| *Flt-1* | Rn00677893_m1 | fms-related tyrosine kinase 4 |
| *VEGFR-2* | Rn00564986_m1 | kinase insert domain receptor |

Quantification was accomplished with the PikoReal Real-Time qPCR System (Applied Biosystems, USA) using TaqMan® Universal PCR Master Mix and universal thermocycling parameters as recommended by Applied Biosystems. RT-PCR samples were run in duplicates using 1 μL cDNA, and the reactions were performed in 96-well plates (Applied Biosystems) in a reaction volume of 20 μL. The gene expression normalization factor for each sample was calculated according to the geometric mean of all three selected reference genes [10]. The Minimum Information for Publication of Quantitative RealTime PCR Experiments (MIQE) guidelines were considered for the performance and interpretations of the qPCR reactions [11]. Gene expression was analyzed according to the relative expression ratio (R-value) obtained from the cycle threshold (CT) value and amplification efficiency (E) value [E = −1+10(−1/slope)]. The expression of targeted genes for each sample provides evidence that these drugs act at the gene level. The entire gene sequence began to amplify at approximately cycle 25 (Fig. 4).

### Statistical analysis

A non-parametric one-way ANOVA test was used to determine the differences in the immunohistochemical expression of Flt-1, Flk-1, and Flt-4 among the experimental groups. Any results where *p* < 0.05 were considered statistically significant. All statistical analyses were conducted using the SPSS software version 21.

## Results

One-way ANOVA revealed a significant difference (*p* < 0.05) for all angiogenic markers (Flt-1, Flk-1, and Flt-4) in all experimental groups.

### Flt-1/VEGFR1

Rapamycin (*M* = 90.1664, *SD* = 7.4487) exhibited better antiangiogenic effects than PF-4 (*M* = 93.7946, *SD* = 7.1303) and rapamycin+PF-4 (*M* = 93.6990, *SD* = 1.8432) (Fig 1) with respect to Flt-1 marker. This demonstrated that PF-4 and rapamycin+PF-4 exerted fewer effects in the suppression of this angiogenesis marker in tumor cells. Contrarily, rapamycin was effective in blocking angiogenic markers through the mTOR signaling pathway.

**Fig 1.**
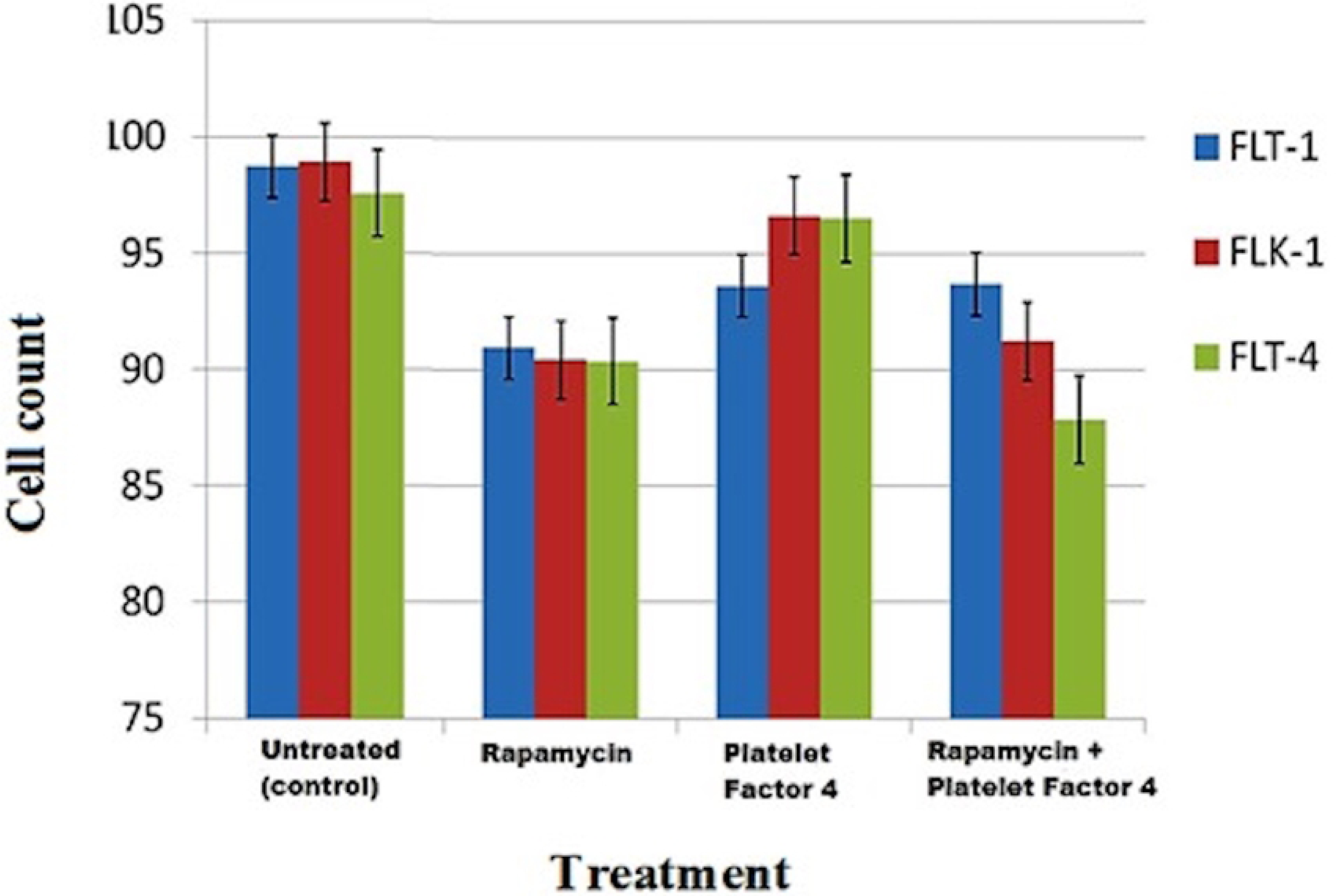
The expression of VEGFRs signaling protein receptor on rat’s mammary carcinoma. The VEGFRs expressions are highly reflected to the efficacy of treatment given to suppress angiogenesis via VEGFRs signaling blockage. All treatment groups showed reduce VEGFRs expressions with *p* value is <0.05.

### Flk-1/VEGFR2

Rapamycin (*M* = 89.9043, *SD* = 7.2542) again revealed better antiangiogenic effects than PF-4 (*M* = 98.9175, *SD* = 2.0487) and rapamycin+PF-4 (*M* = 91.2330, *SD* = 4.0934) in the suppression of the angiogenic marker Flk-1. This marker has the potential to trigger angiogenesis along with other growth factors. These findings were slightly different from that of the Flt-1 marker, as the rapamycin+PF-4 group exhibited better suppression of this marker. The result still demonstrated a lack of synergistic activity between rapamycin and PF-4. Contrarily, PF-4 did not result in good suppression of this receptor, as mentioned in the literature [12].

### Flt-4/VEGFR3

Rapamycin+PF-4 (*M* = 87.8700, *SD* = 4.1620) was very efficacious in suppressing the Flt-4 marker compared with either rapamycin or PF-4 alone. Interestingly, the protein was downregulated more in the combination group than in the other groups, suggesting synergistic activity between these drugs. In contrast, the rapamycin group (*M* = 90.0236, *SD* = 7.2414) exhibited similar effects but to a lesser degree than those of the rapamycin+PF-4 group with respect to Flt-4. Both groups showed significant Flt-4 downregulation. However, PF-4 alone (*M* = 96.5385, *SD* = 1.8304) failed in that the protein was slightly downregulated, but the levels were not significantly different from those after no treatment (control).

### Protein expression and scoring

VEGFRs were highly expressed in the cytoplasm of tumor cells (breast carcinoma) (Figs 2 and 3). The tissues that lacked expression and those without cytoplasmic staining indicated suppressive activity on VEGFRs by the drugs.

**Fig 2.**
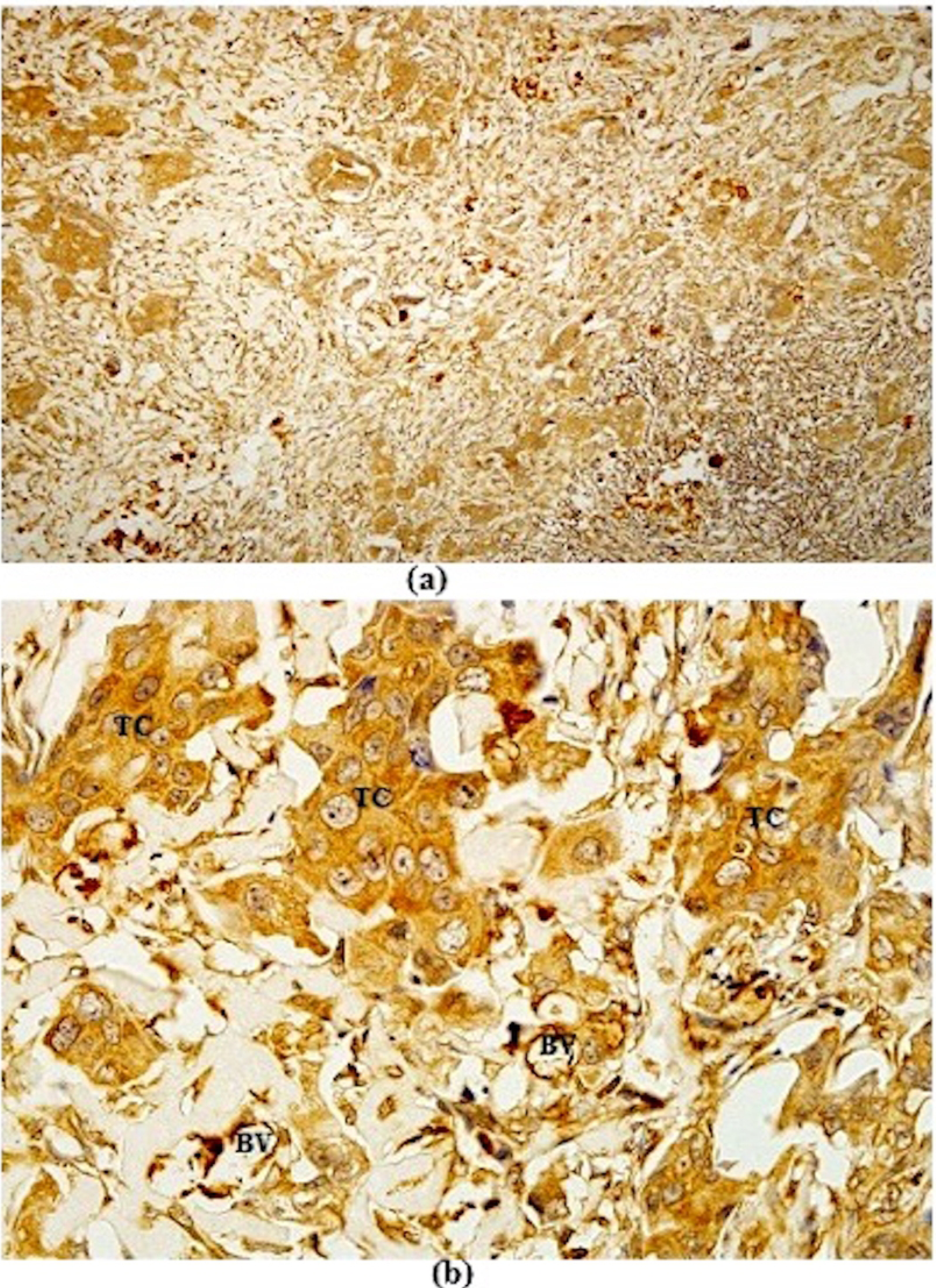
Photomicrograph of NMU-induced rat’s breast tumour tissue. The stain on tumour cells showed the expressions of Flt-1 marker. Flt-1 immunostaining (a) x100 and (b) x400. Tumour cell (TC), blood vessel (BV)

**Fig 3.**
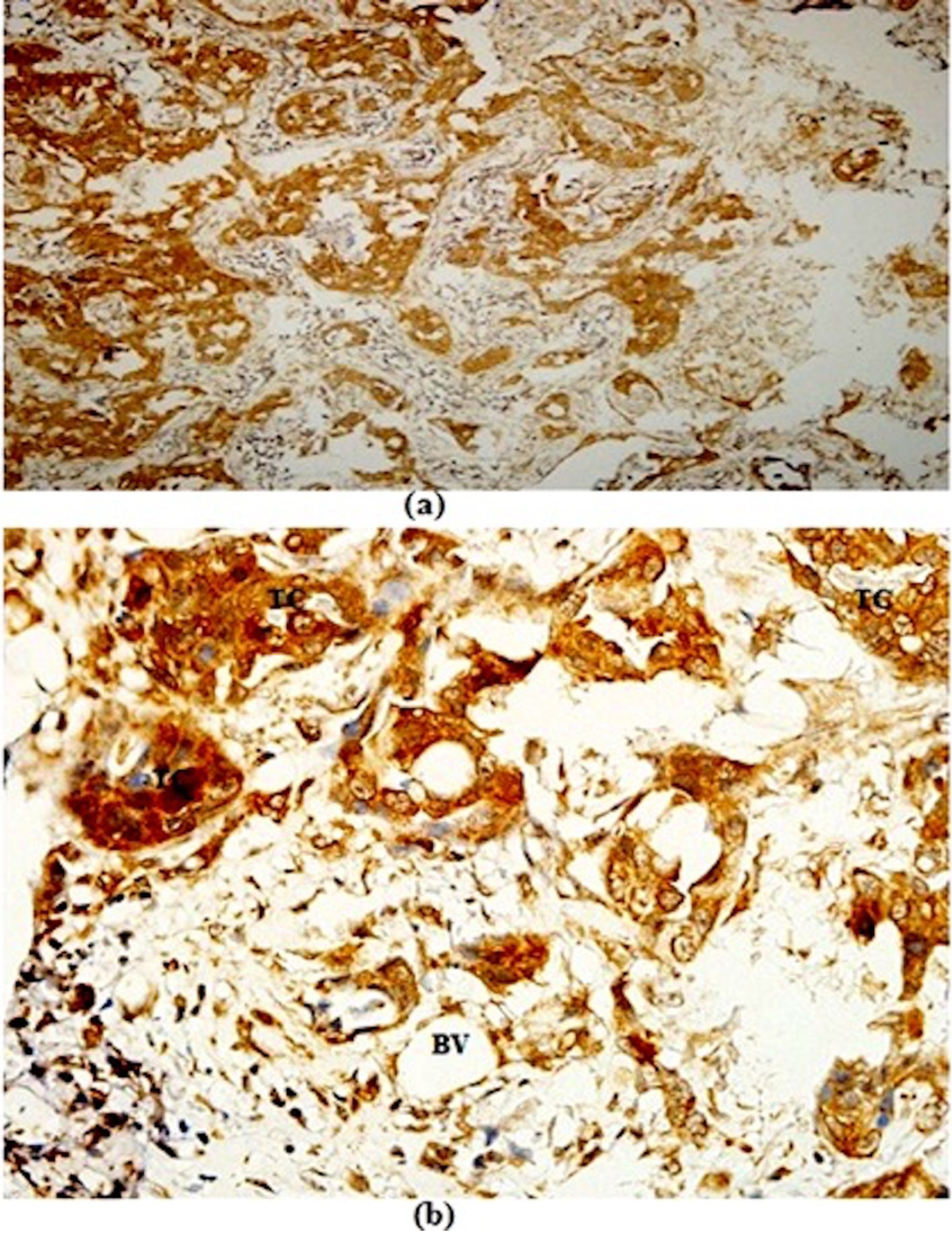
Photomicrograph of NMU-induced rat’s breast tumour tissue. The stain on tumour cells showed the expressions of Flk-1 marker. Flk-1 immunostaining (a) x100 and (b) x400. Tumour cell (TC), blood vessel (BV)

A semiquantitative scoring system was used to determine the protein expression and included counts of positive cells and protein expression intensity. The counting was performed on three selected regions of interest based on the region with the highest protein expressions. The control group had a score of 9 points for both the Flt-1 and Flk-1 markers and a score of 7.5 points for Flt-4, indicating that tumor growth was mostly dependent on the formation of new blood vessels to supply the nutrients needed for that growth. Moreover, the high expression of Flt-4 indicated lymphangiogenesis, which is used by tumor cells as a passageway for metastases.

### Effects of angiogenesis inhibitors

Rapamycin was found to be a good VEGF inhibitor. Based on our findings (Table 2), the angiogenic protein expressions were either completely absent or were expressed at low levels, indicating that rapamycin can substantially downregulate these proteins. This was supported by the semiquantitative scores, where the protein expression decreased from 9 to 6 points. Interestingly, rapamycin+PF-4 exhibited the best inhibitory effects on the Flt-4 ligand, which had a semiquantitative score of 4.5 out of 9 points. We found that the synergism between the drugs was highest for the Flt-4 marker, which plays a significant role in the regulation of lymphatic vessel formation. However, the effects of rapamycin+PF-4 on another two markers, the Flt-1 and Flk-1 ligands, seem less effective, and we found no synergism between the drugs on the suppression of these ligands.

**Table 2.** Semiquantitative score of VEGFR expression in NMU-induced mammary carcinoma in rats. The rapamycin and rapamycin+PF-4 groups exhibited lower scores, whereas the PF-4 group exhibited less-effective suppressive activity on VEGFR markers.

| Treatment | FLT-1 (%) | Scoring | FLK-1 (%) | Scoring | FLT-4 (%) | Scoring |
| --- | --- | --- | --- | --- | --- | --- |
| Control | 98.72 | 9 | 98.92 | 9 | 97.58 | 7.5 |
| Rapamycin | 90.95 | 7.5 | 90.45 | 7.5 | 90.38 | 6 |
| PF-4 | 93.62 | 9 | 96.62 | 7.5 | 96.54 | 7.5 |
| Rapamycin+PF-4 | 93.7 | 7.5 | 91.23 | 7.5 | 87.87 | 4.5 |

### RNA expression

The mRNA expression levels of *Flt-1, Flk-1*, and *Flt-4* genes from the rapamycin, PF-4, and rapamycin+PF-4 treated groups were significantly downregulated compared with the untreated group (Fig 5), with *p-*values of 0.036, 0.018, and 0.000, respectively. Based on these findings, the expression ratio of genes in all intervention groups were downregulated by a mean factor of 0.591 (SE range was 0.441–0.789), 0.458 (SE range was 0.311–0.669), and 0.134 (SE range was 0.078– 0.267) with *p*-values < 0.01. Moreover, we found that the drugs used for intervention had suppressed angiogenesis and lymphangiogenesis at the mRNA level with varying intensities. As demonstrated at the protein level, the administration of rapamycin or rapamycin+PF-4 was found to result in significantly lower than the control (no treatment), whereas intervention with PF-4 led to a lack of suppressive activity on tumor growth indicated by high genes expression, specifically with respect to VEGFRs.

**Fig 4.**
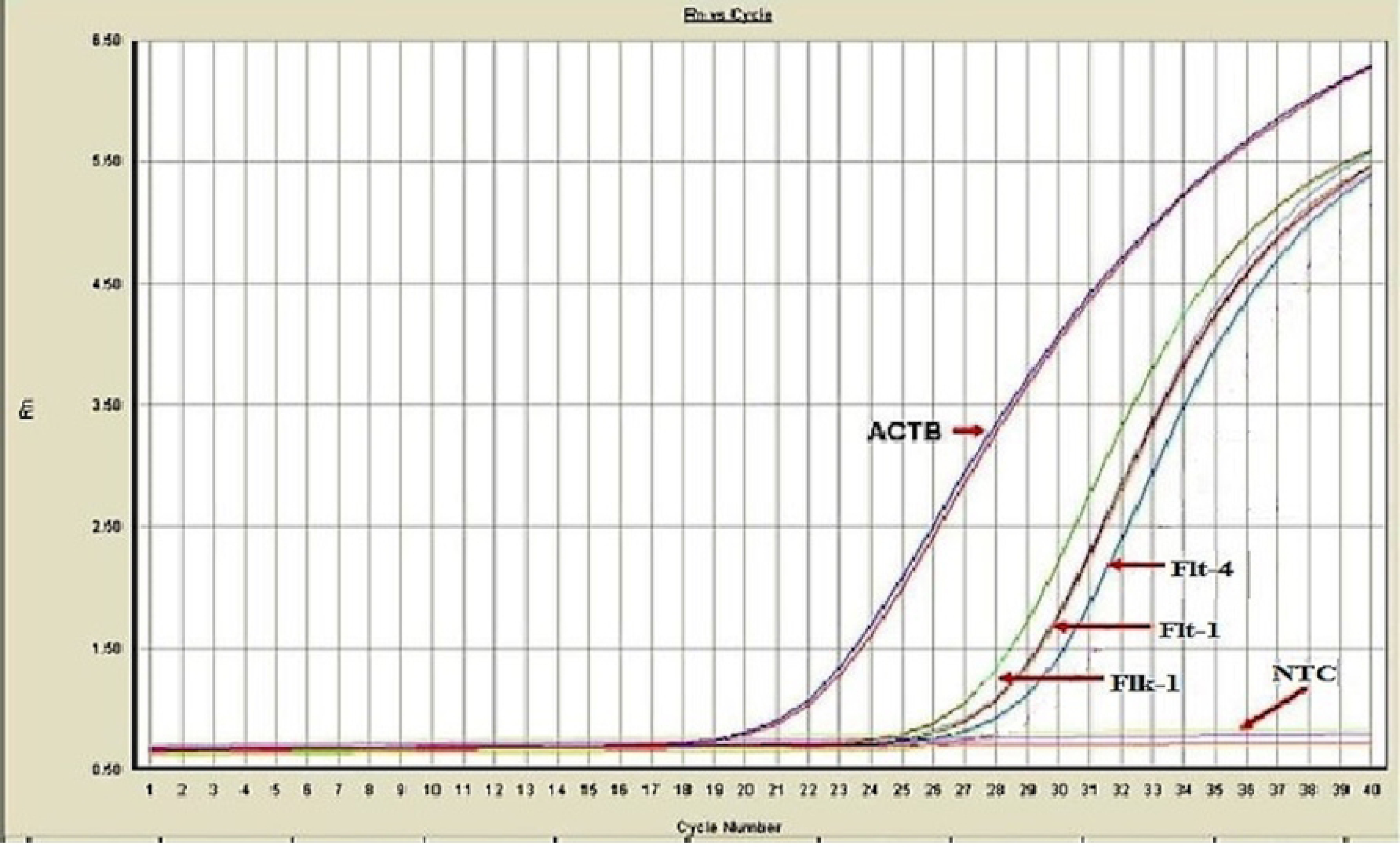

**Fig 5.**
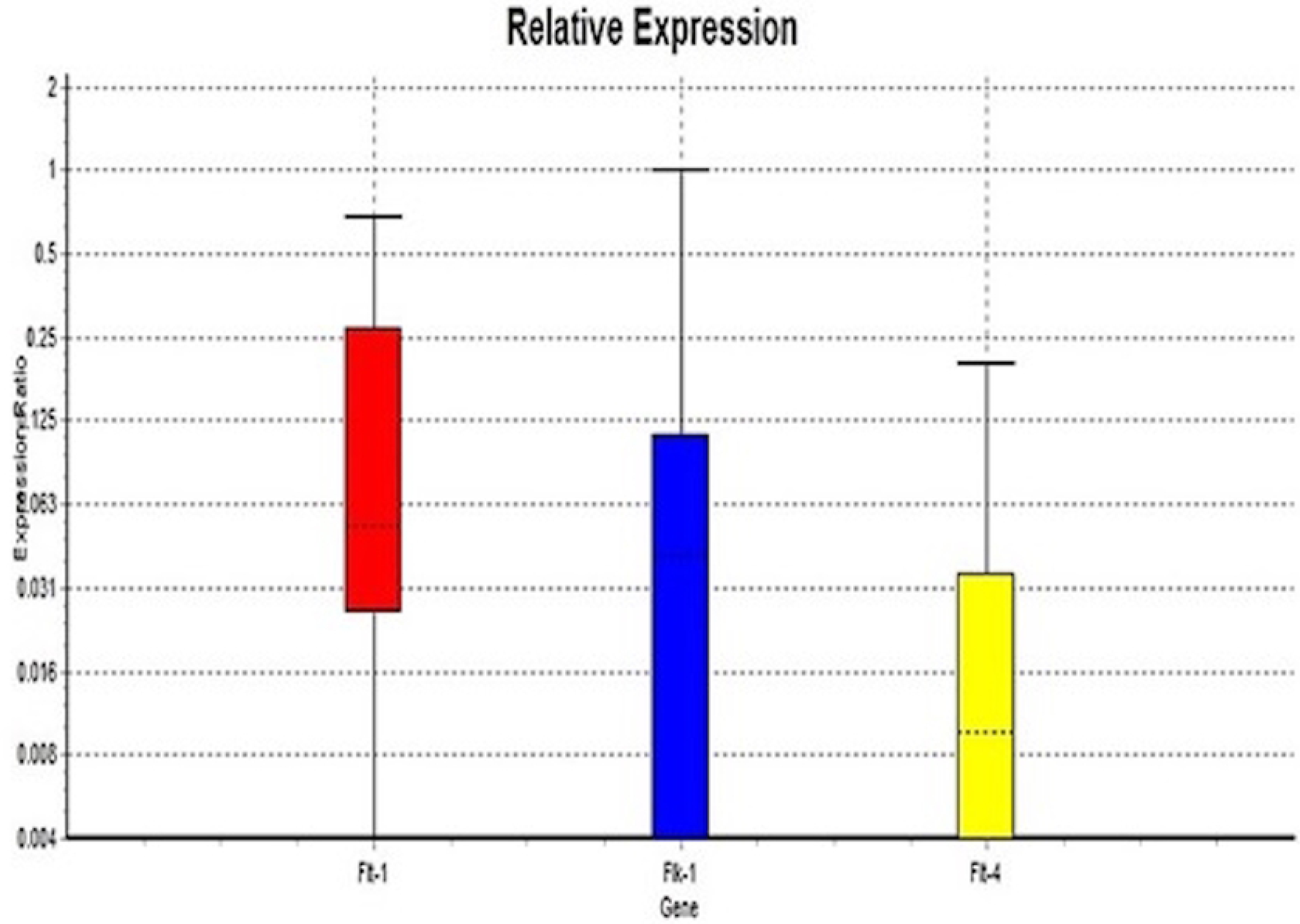
The expression ratio of mRNA angiogenic markers; Flt-1, Flk-1 and Flt-4 on NMU-induced mammary carcinoma under the influence of rapamycin, PF4 and rapamycin+PF4. All VEGFRs marker showed down regulation of gene expression.

## Discussion

In this study, we elucidate the effects of rapamycin and PF-4 as a single dose and in combination on breast carcinoma *in vivo*. We investigated four intervention groups and a control group (untreated). Rapamycin has been used in several medical applications, including as an antifungal agent, an anti-rejection drug after organ transplants, as a treatment for lymphangioleiomyomatosis (LAM), and as an antiproliferative agent (anti-cancer) [13]. Contrarily, PF-4 is a molecule responsible for inducing blood coagulation and activating platelets during platelet aggregation. PF-4 plays a significant role in wound repair and inflammatory reactions [14]. In this study, we observed the relation of synergistic strength between interventional drugs, which are rapamycin and PF-4 against the proliferative and angiogenic activities in breast carcinoma.

Angiogenesis has long been known as fundamental for such diverse physiological processes as embryonic and postnatal development, reproductive functions, and wound repair. The blood vessels provide oxygen and nutrients and carry key regulatory signals to growing tissues [15]. Studies on tumorigenesis have shown that the inflammatory response is sometimes accompanied by increased vascular proliferation. This finding led to the proposal that new blood vessels play a crucial pathogenic role in the regulation of inflammatory processes and to the fulfillment of the demands of proliferating cells. Tumors need nutrients to grow, and their only way to obtain sufficient nutrients is through blood vessels [16]. The mechanisms of the development of new vasculature from the existing vessels require the activation of some upstream signaling receptors, which will then trigger some angiogenic pathways, including the mTOR and PI3K/Akt pathways. Based on these findings, we can hypothesize that aggressive tumor cells express more angiogenic proteins than non-aggressive tumors. This indicates that tumors are highly dependent on angiogenesis, which provides them with additional vessels to fulfill the demands for nutrients and oxygen needed by growing tumor cells. In addition, the vessels also function as a metastatic passageway for tumor cells to spread primarily *via* lymphatic vessels.

Based on the literature, rapamycin acts on mTOR through several upstream pathways. mTOR can be classified into two distinct types, mTOC1 and mTOC2, which are characterized by the presence of raptor and rictor, respectively. Signaling through mTORC1 involves Grb2 and Ras–Raf– MEK–Erk, whereas signaling through mTORC2 involves two different pathways that involve IRS-1 and Grb10 [17]. The VEGF and mTOR pathways are connected by IRS-1 and PI3K/Akt. Inhibition of mTOR pathways will affect the production of VEGF, thus suppressing angiogenesis. In contrast to rapamycin, PF-4 affects the CXCR3 ligand. Activated CXCR3 protein will activate the PI3K/Akt pathway downstream [18]. With the combination of both drugs (rapamycin and PF-4), we hypothesized that they would function synergistically against the PI3K/Akt pathway, which would result in a better prognosis.

This study has noted that after the treatment of rapamycin or rapamycin+PF-4, the expression of VEGFRs was significantly suppressed, whereas PF-4 treatment alone resulted in less suppression of targeted markers. Based on these findings, we report that PF-4 and rapamycin do not function synergistically on VEGFRs, but some evidence suggests that they function antagonistically against the ligands. VEGF-A (Flt-1 and Flk-1) was discovered to be a survival factor for endothelial cells, and it exerts its effects through the PI3K/Akt pathway. High VEGF-A expression in tumor cells suggests that VEGF prevents endothelial cell apoptosis induced by serum starvation.

The expression of angiogenic markers, primarily Flt-1, Flk-1, and Flt-4, was associated with the size and aggressiveness of tumors. Tumor cells in larger, more aggressive tumors require more nutrients and oxygen for their growth [19]. Based on previous studies, Flt-1 protein was found to be highly associated with the production of Flk-1/KDR proteins, which play a significant role in the regulation of angiogenesis and vasculogenesis. Interestingly, with their roles as signaling receptors, Flt-1, Flk-1, and Flt-4 were reported to be responsible for such disorganization and lethality and were found to be negative regulators of VEGF, at least during the early development. In our rat model, Flt-1 was highly expressed in aggressive tumor cells. This indicates that Flt-1 mediates chemotactic signals and potentially extends the role of the receptor, suggesting that this protein promotes mechanisms other than angiogenesis, such as increasing the survival rate of tumor cells, mitogenesis, and permeability-enhancing effects of VEGF rather than being a non-functional or decoy receptor. In the current study, we found that both receptors (Flt-1 and Flk-1) were potentially elevated in epithelial–mesenchymal transition in breast carcinoma and resulted in a rapid proliferation of tumor cells in this tissue.

Angiogenesis highly depends on VEGF protein production, which relies on Flk-1/KDR to activate its pathways [20]. Several previous studies reported that Flk-1 signals act through the PI3K/Akt and PLC-y/Raf-MEK-ERK signaling cascades, which play a significant role in survival, permeability, migration, and proliferation [21]. Bulk production of these proteins is important to sustain a rapidly growing tumor, which requires additional volumes of oxygen and nutrients. In this study, we found that the expression of Flk-1/KDR was prominent, indicating that angiogenesis was very aggressive. The reduced expression of VEGF markers indicates the suppression of angiogenesis.

Flt-4 is the receptor of VEGF-C and VEGF-D and acts as a pivotal regulator of lymphangiogenesis. In this study, we found that the expression of Flt-4 was closely associated with tumor aggressiveness. Previous findings in human breast carcinoma with overexpression of VEGF-C revealed the induction of lymphangiogenesis in and around the tumor. This finding confirms prior evidence that Flt-4 is a vital receptor that regulates the development of new lymphatic vessels, which are an important route for tumor metastases [22]. In this study, we found that Flt-4 was highly expressed in aggressive breast carcinoma but that it was expressed to a lesser degree in the rapamycin- and rapamycin+PF-4-treated groups. These results suggested that both rapamycin and rapamycin+ PF-4 were useful in treating metastatic carcinoma, but that the use of PF-4 alone did not result in good outcomes. Furthermore, we found that the use of antiangiogenic drugs suppressed the development of lymphatic vessels more so than blood vessels. This demonstrates that these drugs have a potent ability to block the interaction between VEGF-C- and Flt-4-soluble VEGFR3 fusion proteins.

## Conclusion

In conclusion, rapamycin is a potent antiangiogenic drug acting on breast carcinoma. The downregulation of VEGFR expression at both protein and mRNA levels in the rapamycin-treated group indicates that rapamycin is best used against an angiogenic stimulant. Conversely, PF-4 has been demonstrated to be a less-effective antiangiogenic drug in this breast carcinoma model.

## Acknowledgments

We would like to acknowledge the Pathology department and animal house staff for their support with laboratory supplies and for imparting some of their wisdom to us. The authors would like to thank Enago (www.enago.com) for the English language review.

## Authors’ contribution

MSMS, HJ, WFWAR, and VG designed experiments, MSMS and TAD performed major experiments and data analysis. MSMS, HJ, and WFWAR performed the statistical analysis. MSMS, HJ, and WFWAR designed the research theme. All authors read and approved the final manuscript.

## Supporting information

Nil

